# Whole-transcriptomic profiling of human cerebral cortex tissues reveals microglia-associated molecular subtypes

**DOI:** 10.1101/2022.05.19.492569

**Authors:** Jiali Zhuang

**Affiliations:** Denovo Biopharma LLC., 10240 Science Center Dr. Ste 120, San Diego, CA, 92121

## Abstract

Microglia is one of the major immune cell types in the human brain and plays pivotal roles in regulating inflammatory and immune response in healthy as well as disease states. By analyzing whole transcriptomic data derived from a large cohort of postmortem cortex tissues, we identified two distinct microglial subtypes within the population. The main difference between the two subtypes lies in the differential expression levels of the C1q complex components, Fc γ receptor (CD16) components and CD14. We validated our discovery in independent cohorts of brain autopsy tissues as well as in RNA-seq data generated from isolated microglia. Future investigations into the causes and physiological implications of these subtypes may shed more light on the homeostasis and regulation of the immune related processes in the brain.

## Introduction

Molecular subtyping is a strategy employed to uncover potential disease heterogeneity in etiology, pathology, progression, and prognosis within patient populations. It has been successfully applied to various types of cancers based on transcriptomic and somatic mutational profiling of tumor tissues (Guinney et al., 2015; Hoadley et al., 2014; Koboldt et al., 2012). The subtype information could inform treatment choice for patients as well as guide the potential development of targeted therapies. The availability of transcriptomic profiling data for biobank scale cohorts has greatly facilitated efforts to identify molecular subtypes for neurodegenerative diseases. For instance, Neff et al. identified three major molecular subtypes of Alzheimer’s Disease (AD) based on their analysis of 1,543 brain tissue transcriptomes across two AD cohorts (Neff et al., 2021). Directly applying clustering algorithms on the whole transcriptomic data often leads to clusters driven largely by variances of cell type compositions within the tissue. An alternative approach is to first identify gene modules that correspond to individual cell types or biological pathways and then interrogate if there exist sample/subject clusters within each gene module based on expression pattern similarity. We applied this strategy to the transcriptomic data of dorsal lateral prefrontal cortex (DLPFC) tissues within the Religious Orders Study-Memory and Aging Project (ROSMAP) cohort and identified two distinct molecular subtypes for microglia-associated genes. Furthermore, we validated our finding in independent cohorts of cortex tissues as well as isolated microglia RNA-sequencing datasets and explored major pathways and processes affected by the subtypes.

## Results

We leveraged whole-transcriptomic dataset derived from a cohort of 629 dorsal lateral prefrontal cortex (DLPFC) autopsy tissues in the ROSMAP study (de Jager et al., 2018) for the discovery of potential molecular subtypes. The cohort includes 249 subjects diagnosed with AD, 168 subjects with minor cognitive impairment (MCI) and 200 age-matched healthy donors with no apparent cognitive impairment. After normalization and batch effect correction (see Materials and Methods), we first performed a high-level decomposition of the aggregated data using Non-negative Matrix Factorization (NMF). Genes enriched in each of the derived components were analyzed to identify common biological processes in Gene Ontology. The top five most enriched processes for each component are listed in Table 1. We noticed that for some components, the enriched processes correspond to the functions or features of certain cell types. For instance, synaptic transmission related processes are the most enriched terms for Component 2 genes, indicating that these genes are preferentially expressed in neurons. Similarly, Component 7 genes seem to be enriched for oligodendrocyte specific processes such as “oligodendrocyte differentiation” and “gliogenesis” (Table 1).

**Table 1.**

|  | GO.ID | Term | Hit | FDR |
| --- | --- | --- | --- | --- |
| Component 1<br>*n* = 263 | GO:0019752 | carboxylic acid metabolic process | 42 | 5.22E-05 |
|  | GO:0006082 | organic acid metabolic process | 44 | 5.22E-05 |
|  | GO:0050877 | nervous system process | 35 | 5.22E-05 |
|  | GO:0055114 | oxidation-reduction process | 39 | 5.22E-05 |
|  | GO:0043436 | oxoacid metabolic process | 43 | 5.22E-05 |
| Component 2<br>*n* = 423 | GO:0007268 | chemical synaptic transmission | 67 | 3.09E-19 |
|  | GO:0098916 | anterograde trans-synaptic signaling | 67 | 3.09E-19 |
|  | GO:0099537 | trans-synaptic signaling | 67 | 5.06E-19 |
|  | GO:0099536 | synaptic signaling | 67 | 6.04E-19 |
|  | GO:0017156 | calcium ion regulated exocytosis | 28 | 8.09E-13 |
| Component 3<br>*n* = 241 | GO:0010817 | regulation of hormone levels | 8 | 1 |
|  | GO:0007186 | G protein-coupled receptor signaling pat... | 8 | 1 |
|  | GO:0008285 | negative regulation of cell proliferatio... | 8 | 1 |
|  | GO:0010648 | negative regulation of cell communicatio... | 12 | 1 |
|  | GO:0023057 | negative regulation of signaling | 12 | 1 |
| Component 4<br>*n* = 45 | GO:0006457 | protein folding | 12 | 2.30E-09 |
|  | GO:0061077 | chaperone-mediated protein folding | 8 | 1.15E-08 |
|  | GO:0009408 | response to heat | 10 | 2.81E-08 |
|  | GO:0006986 | response to unfolded protein | 10 | 4.63E-08 |
|  | GO:1900034 | regulation of cellular response to heat | 8 | 1.12E-07 |
| Component 5<br>*n* = 247 | GO:0006952 | defense response | 82 | <1E-22 |
|  | GO:0002250 | adaptive immune response | 35 | 8.61E-21 |
|  | GO:0006954 | inflammatory response | 45 | 5.62E-20 |
|  | GO:0002252 | immune effector process | 63 | 7.53E-20 |
|  | GO:0034097 | response to cytokine | 63 | 1.59E-19 |
| Component 6<br>*n* = 72 | GO:0007610 | behavior | 10 | 0.17787167 |
|  | GO:0007269 | neurotransmitter secretion | 6 | 0.17787167 |
|  | GO:0099643 | signal release from synapse | 6 | 0.17787167 |
|  | GO:0006836 | neurotransmitter transport | 7 | 0.17787167 |
|  | GO:0099504 | synaptic vesicle cycle | 6 | 0.31829667 |
| Component 7<br>*n* = 361 | GO:0048709 | oligodendrocyte differentiation | 21 | 6.18E-12 |
|  | GO:0042063 | gliogenesis | 27 | 1.78E-08 |
|  | GO:0014003 | oligodendrocyte development | 13 | 2.62E-08 |
|  | GO:0048514 | blood vessel morphogenesis | 36 | 2.62E-08 |
|  | GO:0001525 | angiogenesis | 33 | 2.62E-08 |
| Component 8<br>*n* = 29 | GO:1903708 | positive regulation of hemopoiesis | 6 | 0.00028085 |
|  | GO:0051094 | positive regulation of developmental pro... | 12 | 0.00050553 |
|  | GO:0045597 | positive regulation of cell differentiat... | 10 | 0.00082777 |
|  | GO:0030099 | myeloid cell differentiation | 7 | 0.0009036 |
|  | GO:0002521 | leukocyte differentiation | 7 | 0.0009036 |

To confirm this observation, we conducted analysis on a single cell RNA-seq dataset derived from PFC tissues (Morabito et al., 2020). Clustering of single cells based on the similarity of their gene expression levels revealed distinct cell populations (Figure 1a). By inspecting the expression levels of well-known marker genes across the populations, we were able to assign each population to one of the major cell types in the brain (Figure 1a & 1b). Next, we calculated the average expression levels in each cell population and measured cell type specificity of a gene by comparing its average expression levels across cell populations. We then evaluated the cell type specificity of NMF component enriched genes. Component 1 enriched genes are predominantly astrocyte specific; Component 2 and 6 enriched genes are mostly neuronal genes; Component 7 enriched genes are largely oligodendrocyte specific, whereas Component 5 enriched genes are preferentially expressed in microglia, endothelial cells, and astrocytes (Figure 1c and Supplementary Material Figure 1S).

**Figure 1.**
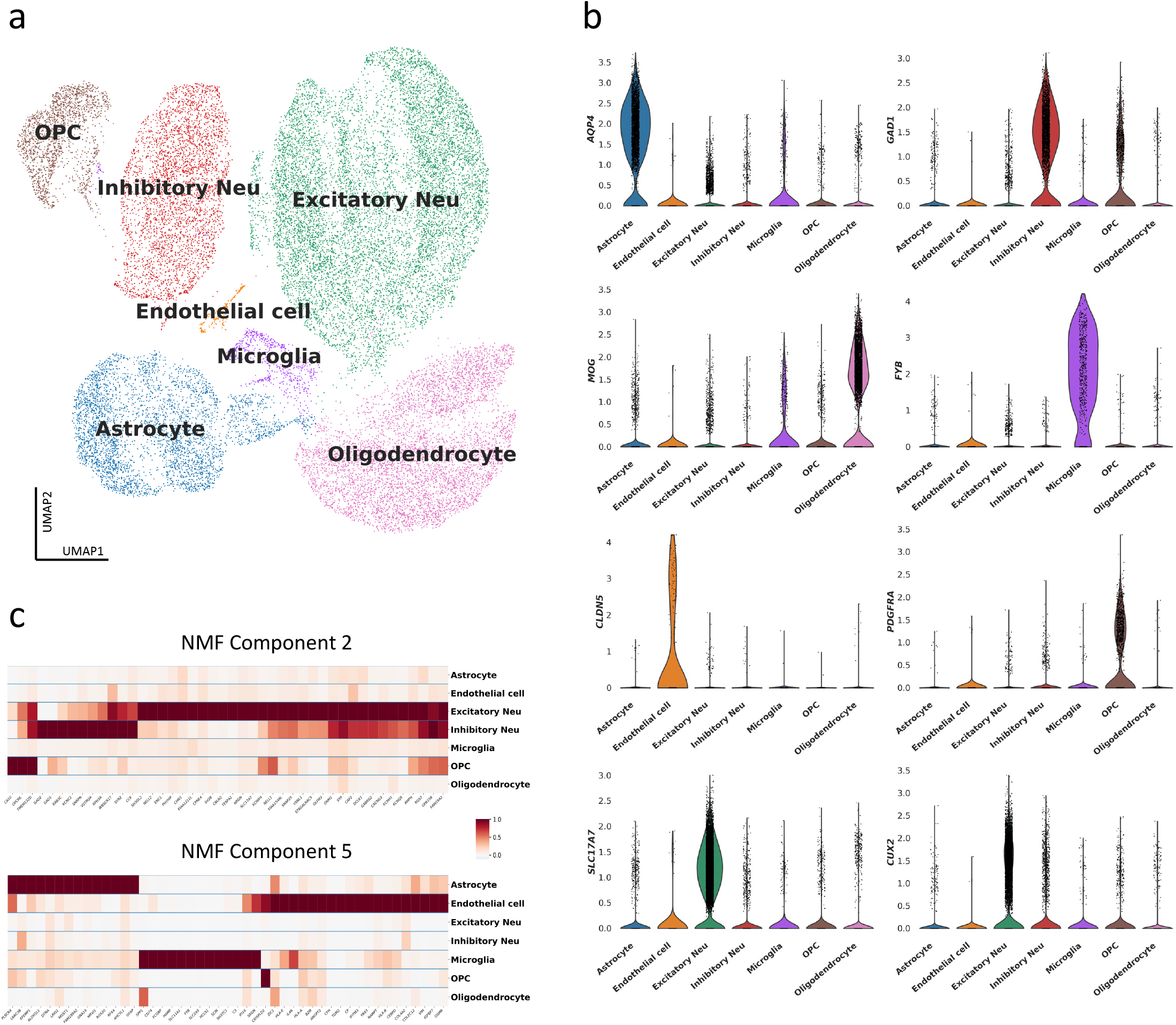
Annotation of NMF components using single-cell brain transcriptomic data. (a) UMAP plot revealing cell populations that represent distinct cell types. (b) Violin plots showing distribution of cell-type specific marker genes across major cell populations. (c) Heatmaps representing cell-type specificity of genes enriched in NMF components 2 and 5. Each column (gene) is normalized to its maximum value.

To obtain a more granular view of the non-neuronal pathways and processes, we employed WGCNA (Langfelder & Horvath, 2008) to group genes enriched in NMF Components 1, 3, 4, 5, 7 and 8 (4,353 non-neuronal genes). This resulted in 18 gene modules (Figure 2a). Gene Ontology enrichment analysis revealed that most modules corresponded to specific biological processes that are essential to the functions and homeostasis of the brain (Supplementary Material Figures S2). For example, “cyan” module genes are enriched for immune response and leucocyte activation related processes; “brown” module genes are enriched for type I interferon signaling pathways and “dark grey” module genes showed enrichment for blood vessel related processes (Figure 2b). Within the “cyan” module genes we recognized several classical markers for microglia cells such as *FYB, INPP5D* and *RUNX1*. Indeed, when inspecting expression levels of the top genes in that module across major cell types using single cell RNA-seq data, we were able to confirm that most “cyan” module genes are specifically expressed in microglia (Figure 2c).

**Figure 2.**
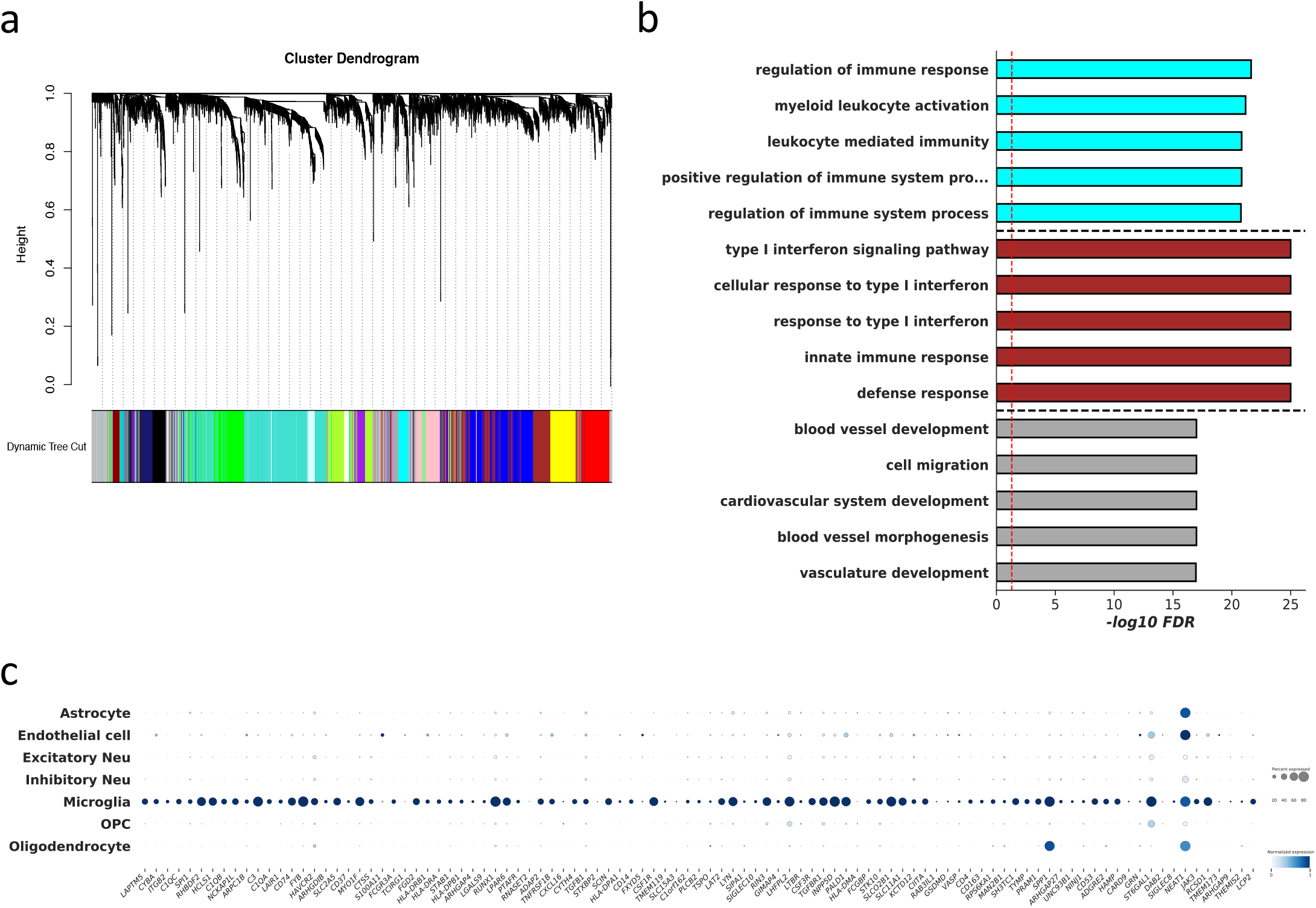
Identification of non-neuronal gene modules with WGCNA. (a) Dendrogram showing gene modules identified with WGCNA. (b) Top 5 Gene Ontology (biological processes) terms for “cyan”, “brown” and “darkgrey” modules. The color of the bars matches the “color” of the respective module. (c) Dotplot showing the cell type specificity of the “cyan” module genes. The colors of the dots denote the average expression level of the genes whereas the sizes of the dots are proportional to the percentage of cells in which the expression of the gene is detected.

Next, we focused on the microglial module (the “cyan” module) genes and investigated whether it is possible to uncover microglia specific subtypes within the ROSMAP cohorts. To this end, we calculated the pairwise Pearson’s correlation coefficients among all samples using the expression levels of the top 100 microglial module genes and performed hierarchical clustering using the pairwise PCC as the distance metric. This analysis revealed two major clusters that included most of the samples (Figure 3a) in the cohort. Samples within the same cluster display higher level of similarity (as manifested by higher PCC) to each other compared with samples from a different cluster. To elucidate the differences between the two clusters and identify the key genes that contribute to such differences, we fitted the expression level data with a linear mixed model (LMM). LMM has the advantage of estimating cluster specific contributions to the coefficients while controlling the variability of overall levels of microglial genes (the intercept) in each donor (see Materials and Methods). We noticed that while for most genes the expression levels are quite comparable for the two clusters, several genes showed considerably elevated expression in one of the two major clusters (Figure 3b). The genes showing the strongest preferential expressions include those encode C1 complement component *C1QA, C1QB* and *C1QC*, Fc γ binding protein *FCGBP*, Fc γ receptor component *FCGR3A* and monocyte differentiation antigen *CD14* (Figure 3b). We validated this discovery in two independent cohorts of cerebral cortex tissue RNA-seq datasets: 258 frontal pole tissues from the Mount Sinai Brain Bank (MSBB) collection (Wang et al., 2018) and 209 frontal cortex (BA 9) tissues from the GTEx consortium (Lonsdale et al., 2013). In both cohorts we found similar patterns of two major clusters whereby one of them is characterized by the elevated levels of C1q component genes, *FCGR3A* and *CD14* (Figure 3c & d, Supplementary Material Figure S3). In fact, when we compared the top ten differential genes between the two clusters in the ROSMAP cohort with those in the MSBB cohort, nine of them are shared between the two cohorts (Supplementary Material Table S1), indicating that the clusters observed in independent cohorts are consistent across independent cohorts and likely represent distinct molecular subtypes.

**Figure 3.**
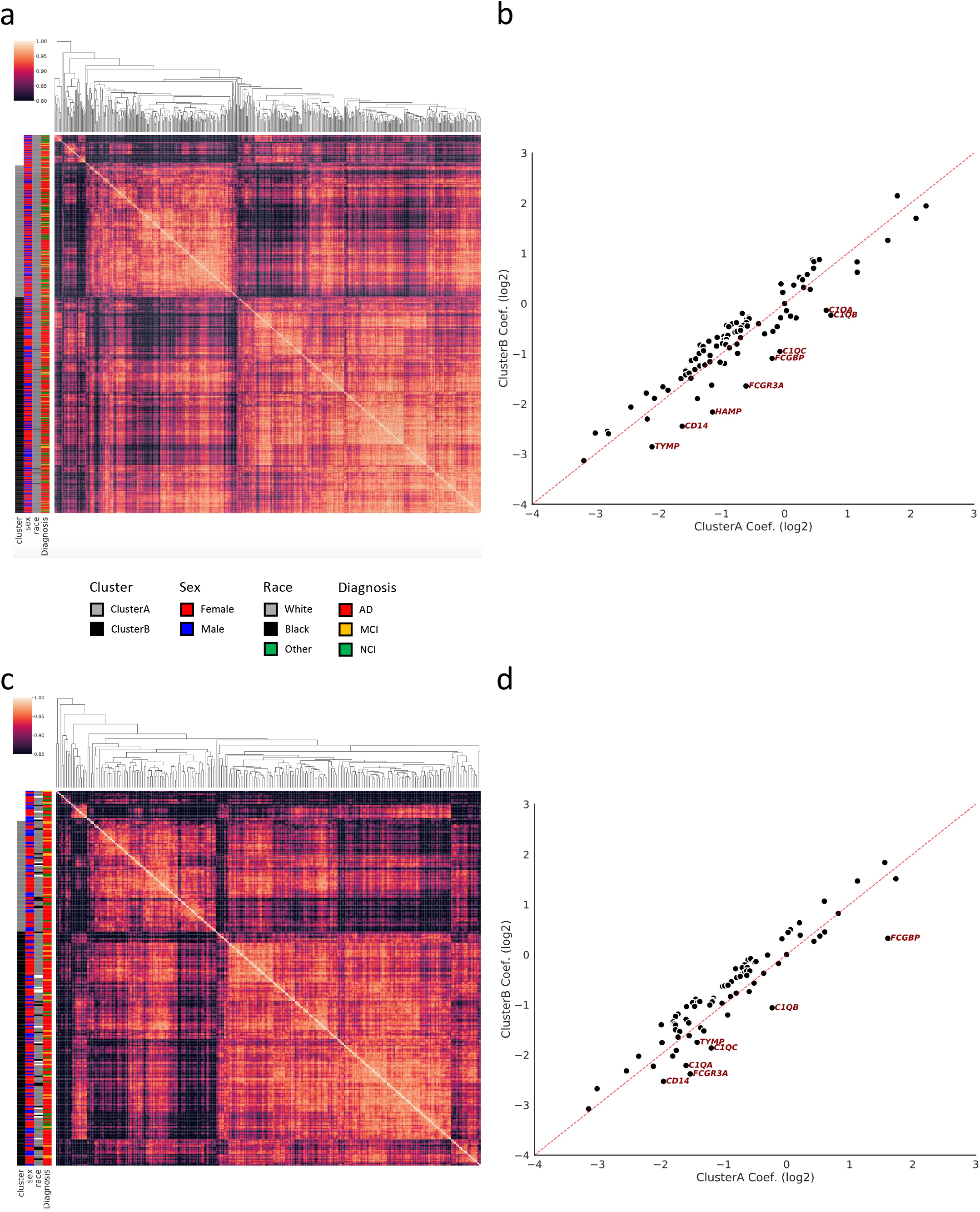
Analysis of microglia gene modules reveals subtypes within large cohorts. (a) & (c) Heatmaps showing subject clusters based on pairwise correlation of microglia genes in ROSMAP (a) and MSBB (c) cohorts. Hierarchical clustering is applied. The color of each grid represents the Person Correlation Coefficient between two donors. The color bars on the left of the heatmap denote cluster membership and demographic traits. (b) & (d) Scatter plots comparing the fitted coefficients of the two clusters in ROSMAP (b) and MSBB (c) cohorts. Labeled genes are the ones showing the highest discrepancies between the two clusters. Dotted red line represents the curve *y* = *x*.

With the potential microglial subtypes identified from transcriptomic profiling of brain tissues, we further inquired whether we would detect similar subtypes from the transcriptomes derived from isolated microglia and if so what pathways and processes are most differentially regulated between the two subtypes. To that end, we leveraged RNA-seq datasets generated from microglia isolated from postmortem autopsy brain tissues from 25 donors of the Netherlands Brain Bank (Alsema et al., 2020). The tissues were dissected from two brain regions: superior parietal lobe (LPS) and superior frontal gyrus (GFS). We first performed hierarchical clustering using the same genes as used in the DLPFC analysis and again identified two major clusters from the microglial RNA-seq data (Figure 4a). Next, we conducted a differential expression analysis comparing the two clusters and found 1,054 differentially expressed genes (FDR < 0.05, Figure 4b). Among these differentially expressed genes are the genes identified from the DLPFC transcriptomes including *C1QA, C1QB, C1QC, FCGR3A* and *CD14* (Figure 4c). Eight out of the ten top differentially expressed genes in DLPFC tissue transcriptomes also show significantly altered expressions (and in the same direction) in the isolated microglial transcriptomes, which is significantly higher than expected by chance (p-value = 1.36e-5, hypergeometric test). This indicated that the microglial subtypes we discovered from DLPFC tissue RNA-seq dataset is replicated in isolated microglia. Brain tissue transcriptomes contain transcripts from various cell types and therefore we can only focus on those genes specifically expressed in microglia (the “cyan” module). RNA-seq dataset derived from isolated microglia, on the other hand, does not have such limitation and allows us to compare the two subtypes in a more comprehensive manner. We performed KEGG and Reactome pathways enrichment analysis for the differentially expressed genes using pathfindR (Ulgen et al., 2019). Among the enriched KEGG pathways are TNF signaling pathway and Fc γ receptor-mediated phagocytosis, while interleukin signaling pathway and TLR4-related signaling pathways are notable among the enriched Reactome pathways (Supplementary Material Figure S4).

**Figure 4.**
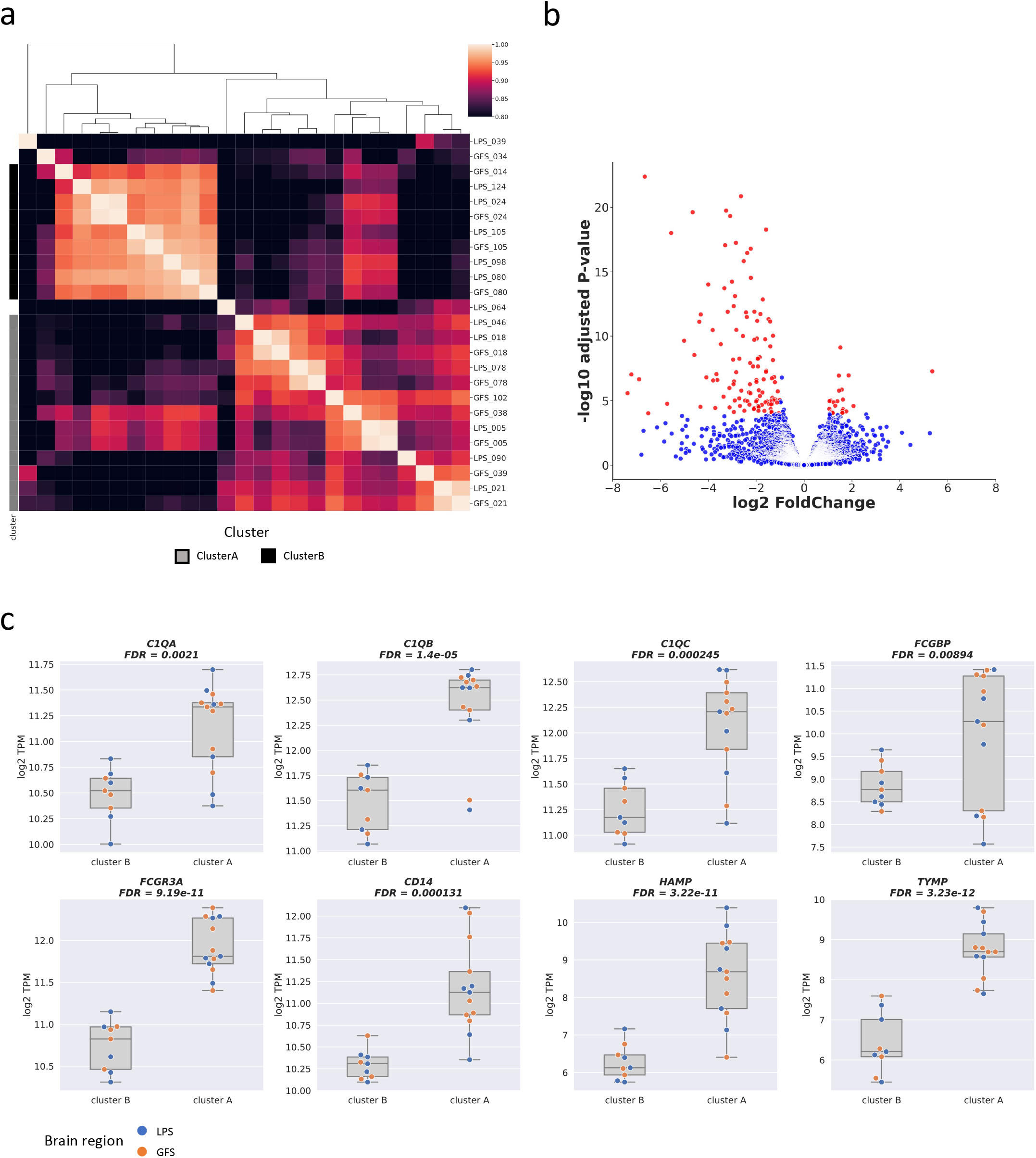
Similar subtypes are observed in microglia RNA-seq dataset. (a) Heatmap showing hierarchical clustering based on microglia RNA-seq data. The color of each grid represents the Person Correlation Coefficient between two samples. The color bars on the left of the heatmap denote cluster membership. (b) Volcano plot showing differentially expressed genes between the two clusters. Genes that are up- or down-regulated by more than 2-fold and have adjusted p-values smaller than 1e-4 are colored in red. (c) Boxplots comparing the expression levels of *C1QA, C1QB, C1QC, FCGBP, FCGR3A, CD14, HAMP* and *TYMP* in microglia between the two clusters.

A key question regarding the microglial subtypes we discovered is if there is any clinical or physiological metrics associated with it. For donors within the ROSMAP study, several demographic and clinical information are available. Since the design of the ROSMAP study has a particular focus on Alzheimer’s Disease, the records include the AD diagnostic status, APOE genotype, Braak staging and CERAD score of the donors. We tested the association between the microglial subtypes and the aforementioned clinical measurements as well as sex of the subjects, RNA integrity number (RIN) and postmortem interval of the samples (PMI). After correcting for multiple testing, none of the characteristics we analyzed was significantly associated with microglial subtypes (Figure 5a, False Discovery Rate < 0.05). We further investigated whether there are any genomic markers associated with the microglial subtypes. The ROSMAP consortium conducted whole genome sequencing on 552 of the 629 donors included in our analysis, allowing us to use the genotype information for a genome-wide association analysis. We tested ∼6.28 million variants with minor allele frequency greater than 5% and did not detect any variants with significant association (using threshold p-value = 5e-8, Figure 5b & c). This indicated that either genetic factors did not play a major role in the determination of the microglial subtypes, or our current cohort lacked sufficient power to detect the effects.

**Figure 5.**
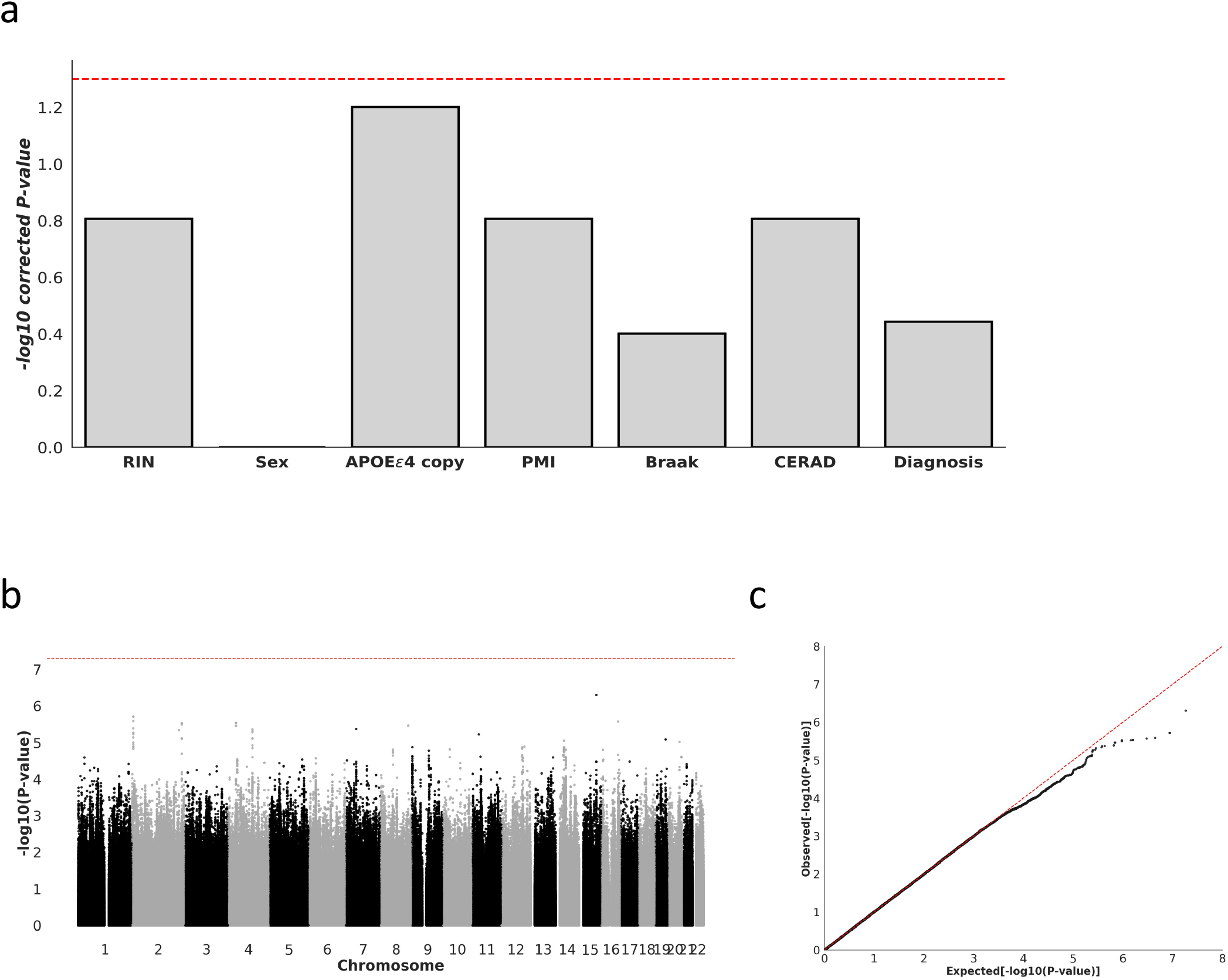
The microglia subtypes are not explained by any of the sample characteristics, patient demographic/pathological traits or genomic variants we investigated. (a) Bar plots showing the adjusted p-values of association between various clinical/demographical features and the microglia subtype. (b) & (c) Manhattan plot (b) and Quantile-Quantile plot (c) of genome-wide association study analysis between genomic variants and microglia subtypes. Red dotted lines in (a) and (b) represent levels of statistical significance.

## Discussion

Transcriptomic data generated from postmortem brain tissue RNA sequencing typically reflects contributions from multiple cell types and pathways. The ability to deconvolute and dissect the data computationally allows us to individually evaluate cell types or pathways of interest. The availability of single cell RNA-seq data supports and facilitates the interpretation and annotation of the results. In this study, we employed a computational workflow that combines NMF and WGCNA to deconvolute the transcriptomic data of 629 DLPFC autopsy tissues in the ROSMAP study and identified gene modules that correspond to specific cell types and biological processes in the central nervous system. By focusing on the gene module associated with the cell type or pathway of interest, we were able to target our analysis on that particular cell type or pathway even though the transcriptomic data includes information from many other cell types or pathways. This mode of investigation is useful considering that cell sorting and enrichment are often laborious and time-consuming for a large cohort of tissues, and it allows us to directly harness the existing high-quality tissue RNA-seq datasets for the investigation of specific cell types or pathways without requiring cell sorting.

When analyzing the expression profiles of microglial module genes within the ROSMAP cohort, we discovered two distinct subtypes characterized by the differential expression of a small number of genes. This observation is replicated in two additional independent cohorts, suggesting that it likely reflects a genuine feature within the human population. Most of the genes differentially expressed between the two subtypes are related to the recognition and clearance of invading foreign microbes. For instance, *C1QA, C1QB* and *C1QC* encode components of the C1 complex in the complement system, which is responsible for binding to the antigen-antibody complexes and initiating attacks on the membrane of the microbes (Reid, 2018). *FCGR3A* encodes a component of the Fc fragment of IgG receptor III (also known as CD16), which plays key roles in the removal of antigen-antibody complexes and mediating antibody dependent cellular cytotoxicity (Mahaweni et al., 2018). Cluster of differentiation 14 (*CD14*) encodes a protein that (with the help of other proteins) binds to bacterial lipopolysaccharide (LPS) as well as other pathogen-associated molecular patterns and activates Toll-like receptor 4 (TLR4) along with its downstream signaling pathway (Zanoni et al., 2011). To obtain a more comprehensive picture of the pathways driving the inter-subtypes differences, we searched for pathways that are enriched in genes differentially expressed between the two subtypes. Among the enriched KEGG pathways are “TNF signaling pathway”, which is a key regulator of inflammation; and “Fc γ receptor mediated phagocytosis”, which includes *FCGR3A*. The enriched Reactome pathways include TLR4 downstream processes such as “MyD88-independent TLR4 cascade” and “TRIF-mediated TLR4 signaling”, which are activated by the presence of invading pathogens; and interleukins signaling pathways which are activated upon the triggering of TLR4 signaling pathway. These findings imply that the main differences between the transcriptomes of the two microglial subtypes might be related to immune responses to microbial pathogens in the brain.

Our analysis revealed that the microglial subtypes are not significantly associated with the onset or progression of Alzheimer’s Disease within our study cohort. Therefore, the physiological and pathological implications of the cortex microglial subtype remain to be discovered. Larger cohorts with more diverse backgrounds and involving more disease phenotypes will likely be needed to solve the puzzle. Another potential usage of the molecular subtype is to predict response to novel therapies, especially those targeting the inflammatory and immune response pathways in the brain. Since microglia cells are a critical component of the CNS immune system, it is possible that microglial subtypes might influence the efficacy of immune-modulating interventions. We uncovered the microglial subtypes by analyzing the transcriptomic data of the postmortem cortex tissues, but brain tissues are unavailable for studies involving living subjects. Therefore, we attempted to find genetic variants whose genotypes strongly associate with the molecular subtypes and can be used as surrogates to predict subtype status of the subject without requiring brain biopsy. With the genome-wide association analysis conducted on the ROSMAP cohort, we unfortunately did not find any variant that significantly associates with the microglial subtypes. Future studies with increased power or testing alternative surrogates are needed in the search for more efficient approaches to determine the microglial subtype status.

## Materials and Methods

### RNA-seq data processing

RNA-seq and genomic sequencing data for the ROSMAP and MSBB studies are hosted by the AMP-AD consortium and is accessible through the Synapse portal (https://www.synapse.org/#!Synapse:syn2580853/wiki/409840). Demographic and clinical information are available through the same portal (syn3191087 for ROSMAP; syn6101474 for MSBB). Processed and harmonized read-count matrix tables were directly downloaded from the Synapse portal (syn10507739 for ROSMAP; syn10507730 for MSBB). Briefly, reads were aligned to the GENCODE24 (GRCh38) reference genome using STAR, with “twopassMode” set as Basic. Transcript abundances were estimated for each sample using Sailfish. For more details about RNA-seq data processing please refer to https://www.synapse.org/#!Synapse:syn17010685. RNA-seq data of isolated microglia from the Netherlands Brain Bank study was downloaded from Gene Expression Omnibus (GSE146639). In this study, we analyzed the RNA-seq data at the gene level by summing up read counts over all isoforms belonging to the same gene. Genes that were detected (>= 5 counts per million) in less than 75% of the samples were excluded from subsequent analysis. The “removeBatchEffect” function in the “limma” R library (version 3.42.2) (Ritchie et al., 2015) was used to correct the effects of sequencing batch and RNA Integrity Number (RIN) after regression using voom (Law et al., 2014). Variant calling results from whole genome sequencing for ROSMAP study were downloaded from Synapse portal (syn11707420). Single-cell RNA-seq data was downloaded from Synapse portal (syn18915937). The processing of single-cell RNA-seq data including filtering, transformation, normalization, dimension reduction and visualization was implemented using Scanpy (version 1.8.2) library in Python (Wolf et al., 2018). Differential gene expression analysis was performed using DESeq2 (version 1.26.0) R package (Love et al., 2014).

### Non-negative Matrix Factorization and gene module analysis

NMF decomposition was implemented in Python using the “NMF” function of the scikit learn library (version 0.23.2) (Pedregosa FABIANPEDREGOSA et al., 2011) with number of components set to eight (n_components = 8), maximum number of iterations set to 1,000,000 (max_iter = 1000000) and no regularization penalty (alpha = 0). The mean and standard deviation of the read counts were calculated for each gene and capped at 5 standard deviations. Before the decomposition, the read counts for each gene were normalized to the 3rd quartile (75th percentile) to ensure that genes are comparable. The resulting loading matrix was normalized by the sum of weights for each gene to determine the relative contributions to each component. For a gene-component pair if the relative contribution by the gene to the component is greater than 35%, the gene is considered “enriched” in that component. Gene module analysis was implemented using the “WGCNA” (version 1.70-3) package in R (Langfelder & Horvath, 2008). In the construction of adjacency matrix, the soft thresholding power set to 10. After gene clustering, we merged those gene modules whose eigengenes have correlation greater then 0.75.

### Sample clustering based on microglial genes

The 100 genes with the highest contribution to the eigengene of the microglial module (“cyan” module) were selected for subsequent analysis. Hierarchical clustering was performed on the expression levels of those 100 genes across the donors using the “linkage” function in the Scipy (version 1.6.1) (Virtanen et al., 2020) library (scipy.cluster.hierarchy) in Python with metric set to “correlation” and method set to “average”. Linear Mixed Model (LMM) was used to ascertain the average expression profiles of the two major clusters while controlling for individual variations and was implemented in Python with the statsmodels.formula.api.mixedlm function in the statsmodels library (version 0.12.2) (Perktold & Seabold, 2010). The formula for the mixed model is *E*_*ijk*_ = *g*_*i*_ + *c*_*j*_ + *a*_*ij*_ + *σ*_*k*_ + *ε*_*ijk*_, where *E*_*ijk*_ stands for the expression level of gene *i* in donor *k* belonging to cluster *j, g*_*i*_, the coefficient for gene *i, c*_*j*_ the coefficient for cluster *j, a*_*ij*_, the effects of membership in cluster *j* on gene *i, σ*_*k*_ the intercept for donor *k* and *ε*_*ijk*_the noise. *σ*_*k*_ is treated as the random effect in the model while *g*_*i*_,*c*_*j*_, and *a*_*ij*_ are treated as the fixed effects. The fitting of the LMM is performed after logarithm (log2) transformation.

### Association of microglial subtypes with demographic/clinical traits and genomic variants

The association of microglial subtype with categorical traits such as sex and AD status are measured by Fisher’s Exact test with the “scipy.stats. fisher_exact” function in the Scipy (version 1.6.1) (Virtanen et al., 2020) library. The association with numerical traits including RIN, PMI, APOE ε4 allele copy number, Braak staging and CERAD score are calculated by comparing the values of those traits of the two subtypes using Mann-Whitney test (scipy.stats.mannwhitneyu). The association with the genomic variants was performed using PLINK (version v1.90b6.17) (Purcell et al., 2007) with the logistic regression mode (--logistic -- geno --maf 0.05 --hwe 0.000001) and treating sex, age, and PMI as covariates.

### Gene Ontology and pathway enrichment analysis

Enrichment for Gene Ontology biological process terms was performed using the “topGO” package (version 2.38.1) in R. Benjamini-Hochberg method was used to correct for multiple testing. Pathway enrichment analysis was performed using pathfindeR (version 1.4.1) (Ulgen et al., 2019) with options “min_gset_size=50, max_gset_size=500”.

## Acknowledgement

The results published here are in part based on data obtained from the AD Knowledge Portal (https://adknowledgeportal.org). Data generation was supported by the following NIH grants: P30AG10161, P30AG72975, R01AG15819, R01AG17917, R01AG036836, U01AG46152, U01AG61356, U01AG046139, P50 AG016574, R01 AG032990, U01AG046139, R01AG018023, U01AG006576, U01AG006786, R01AG025711, R01AG017216, R01AG003949, R01NS080820, U24NS072026, P30AG19610, U01AG046170, RF1AG057440, and U24AG061340, and the Cure PSP, Mayo and Michael J Fox foundations, Arizona Department of Health Services and the Arizona Biomedical Research Commission. We thank the participants of the Religious Order Study and Memory and Aging projects for the generous donation, the Sun Health Research Institute Brain and Body Donation Program, the Mayo Clinic Brain Bank, and the Mount Sinai/JJ Peters VA Medical Center NIH Brain and Tissue Repository. Data and analysis contributing investigators include Nilüfer Ertekin-Taner, Steven Younkin (Mayo Clinic, Jacksonville, FL), Todd Golde (University of Florida), Nathan Price (Institute for Systems Biology), David Bennett, Christopher Gaiteri (Rush University), Philip De Jager (Columbia University), Bin Zhang, Eric Schadt, Michelle Ehrlich, Vahram Haroutunian, Sam Gandy (Icahn School of Medicine at Mount Sinai), Koichi Iijima (National Center for Geriatrics and Gerontology, Japan), Scott Noggle (New York Stem Cell Foundation), Lara Mangravite (Sage Bionetworks).

## Reference

Alsema, A. M., Jiang, Q., Kracht, L., Gerrits, E., Dubbelaar, M. L., Miedema, A., Brouwer, N., Hol, E. M., Middeldorp, J., van Dijk, R., Woodbury, M., Wachter, A., Xi, S., Möller, T., Biber, K. P., Kooistra, S. M., Boddeke, E. W. G. M., & Eggen, B. J. L. (2020). Profiling Microglia From Alzheimer’s Disease Donors and Non-demented Elderly in Acute Human Postmortem Cortical Tissue. Frontiers in Molecular Neuroscience, 13. https://doi.org/10.3389/fnmol.2020.00134

de Jager, P. L., Ma, Y., McCabe, C., Xu, J., Vardarajan, B. N., Felsky, D., Klein, H. U., White, C. C., Peters, M. A., Lodgson, B., Nejad, P., Tang, A., Mangravite, L. M., Yu, L., Gaiteri, C., Mostafavi, S., Schneider, J. A., & Bennett, D. A. (2018). Data descriptor: A multi-omic atlas of the human frontal cortex for aging and Alzheimer’s disease research. Scientific Data, 5. https://doi.org/10.1038/sdata.2018.142

Guinney, J., Dienstmann, R., Wang, X., de Reyniès, A., Schlicker, A., Soneson, C., Marisa, L., Roepman, P., Nyamundanda, G., Angelino, P., Bot, B. M., Morris, J. S., Simon, I. M., Gerster, S., Fessler, E., de Sousa. E Melo, F., Missiaglia, E., Ramay, H., Barras, D., … Tejpar, S. (2015). The consensus molecular subtypes of colorectal cancer. Nature Medicine, 21(11), 1350–1356. https://doi.org/10.1038/nm.3967

Hoadley, K. A., Yau, C., Wolf, D. M., Cherniack, A. D., Tamborero, D., Ng, S., Leiserson, M. D. M., Niu, B., McLellan, M. D., Uzunangelov, V., Zhang, J., Kandoth, C., Akbani, R., Shen, H., Omberg, L., Chu, A., Margolin, A. A., Van’t Veer, L. J., Lopez-Bigas, N., … Zou, L. (2014). Multiplatform analysis of 12 cancer types reveals molecular classification within and across tissues of origin. Cell, 158(4), 929–944. https://doi.org/10.1016/j.cell.2014.06.049

Koboldt, D. C., Fulton, R. S., McLellan, M. D., Schmidt, H., Kalicki-Veizer, J., McMichael, J. F., Fulton, L. L., Dooling, D. J., Ding, L., Mardis, E. R., Wilson, R. K., Ally, A., Balasundaram, M., Butterfield, Y. S. N., Carlsen, R., Carter, C., Chu, A., Chuah, E., Chun, H. J. E., … Palchik, J. D. (2012). Comprehensive molecular portraits of human breast tumours. Nature, 490(7418), 61–70. https://doi.org/10.1038/nature11412

Langfelder, P., & Horvath, S. (2008). WGCNA: An R package for weighted correlation network analysis. BMC Bioinformatics, 9. https://doi.org/10.1186/1471-2105-9-559

Law, C. W., Chen, Y., Shi, W., & Smyth, G. K. (2014). voom: precision weights unlock linear model analysis tools for RNA-seq read counts. In Genome Biology (Vol. 15). http://genomebiology.com/2014/15/2/R29

Lonsdale, J., Thomas, J., Salvatore, M., Phillips, R., Lo, E., Shad, S., Hasz, R., Walters, G., Garcia, F., Young, N., Foster, B., Moser, M., Karasik, E., Gillard, B., Ramsey, K., Sullivan, S., Bridge, J., Magazine, H., Syron, J., … Moore, H. F. (2013). The Genotype-Tissue Expression (GTEx) project. In Nature Genetics (Vol. 45, Issue 6, pp. 580–585). https://doi.org/10.1038/ng.2653

Love, M. I., Huber, W., & Anders, S. (2014). Moderated estimation of fold change and dispersion for RNA-seq data with DESeq2. Genome Biology, 15(12). https://doi.org/10.1186/s13059-014-0550-8

Mahaweni, N. M., Olieslagers, T. I., Rivas, I. O., Molenbroeck, S. J. J., Groeneweg, M., Bos, G. M. J., Tilanus, M. G. J., Voorter, C. E. M., & Wieten, L. (2018). A comprehensive overview of FCGR3A gene variability by full-length gene sequencing including the identification of V158F polymorphism. Scientific Reports, 8(1). https://doi.org/10.1038/s41598-018-34258-1

Morabito, S., Miyoshi, E., Michael, N., & Swarup, V. (2020). Integrative genomics approach identifies conserved transcriptomic networks in Alzheimer’s disease. Human Molecular Genetics, 29(17), 2899–2919. https://doi.org/10.1093/hmg/ddaa182

Neff, R. A., Wang, M., Vatansever, S., Guo, L., Ming, C., Wang, Q., Wang, E., Horgusluoglu-Moloch, E., Song, W.-M., Li, A., Castranio, E. L., Tcw, J., Ho, L., Goate, A., Fossati, V., Noggle, S., Gandy, S., Ehrlich, M. E., Katsel, P., … Zhang, B. (2021). Molecular subtyping of Alzheimer’s disease using RNA sequencing data reveals novel mechanisms and targets. In Sci. Adv (Vol. 7, Issue 6). http://advances.sciencemag.org/

Pedregosa Fabianpedregosa, F., Michel, V., Grisel Oliviergrisel, O., Blondel, M., Prettenhofer, P., Weiss, R., Vanderplas, J., Cournapeau, D., Pedregosa, F., Varoquaux, G., Gramfort, A., Thirion, B., Grisel, O., Dubourg, V., Passos, A., Brucher, M., Perrot andÉdouardand, M., Duchesnay, andÉdouard, & Duchesnay Edouardduchesnay Fré., (2011). Scikit-learn: Machine Learning in Python Gaël Varoquaux Bertrand Thirion Vincent Dubourg Alexandre Passos PEDREGOSA, VAROQUAUX, GRAMFORT ET AL. Matthieu Perrot. In Journal of Machine Learning Research (Vol. 12). http://scikit-learn.sourceforge.net.

Perktold, J., & Seabold, S. (2010). Statsmodels: Econometric and Statistical Modeling with Python Quantitative histology of aorta View project Statsmodels: Econometric and Statistical Modeling with Python. https://www.researchgate.net/publication/264891066

Purcell, S., Neale, B., Todd-Brown, K., Thomas, L., Ferreira, M. A. R., Bender, D., Maller, J., Sklar, P., de Bakker, P. I. W., Daly, M. J., & Sham, P. C. (2007). PLINK: A tool set for whole-genome association and population-based linkage analyses. American Journal of Human Genetics, 81(3), 559–575. https://doi.org/10.1086/519795

Reid, K. B. M. (2018). Complement component C1q: Historical perspective of a functionally versatile, and structurally unusual, serum protein. Frontiers in Immunology, 9(APR). https://doi.org/10.3389/fimmu.2018.00764

Ritchie, M. E., Phipson, B., Wu, D., Hu, Y., Law, C. W., Shi, W., & Smyth, G. K. (2015). Limma powers differential expression analyses for RNA-sequencing and microarray studies. Nucleic Acids Research, 43(7), e47. https://doi.org/10.1093/nar/gkv007

Ulgen, E., Ozisik, O., & Sezerman, O. U. (2019). PathfindR: An R package for comprehensive identification of enriched pathways in omics data through active subnetworks. Frontiers in Genetics, 10(SEP). https://doi.org/10.3389/fgene.2019.00858

Virtanen, P., Gommers, R., Oliphant, T. E., Haberland, M., Reddy, T., Cournapeau, D., Burovski, E., Peterson, P., Weckesser, W., Bright, J., van der Walt, S. J., Brett, M., Wilson, J., Millman, K. J., Mayorov, N., Nelson, A. R. J., Jones, E., Kern, R., Larson, E., … Vázquez-Baeza, Y. (2020). SciPy 1.0: fundamental algorithms for scientific computing in Python. Nature Methods, 17(3), 261–272. https://doi.org/10.1038/s41592-019-0686-2

Wang, M., Beckmann, N. D., Roussos, P., Wang, E., Zhou, X., Wang, Q., Ming, C., Neff, R., Ma, W., Fullard, J. F., Hauberg, M. E., Bendl, J., Peters, M. A., Logsdon, B., Wang, P., Mahajan, M., Mangravite, L. M., Dammer, E. B., Duong, D. M., … Zhang, B. (2018). The Mount Sinai cohort of large-scale genomic, transcriptomic and proteomic data in Alzheimer’s disease. Scientific Data, 5. https://doi.org/10.1038/sdata.2018.185

Wolf, F. A., Angerer, P., & Theis, F. J. (2018). SCANPY: Large-scale single-cell gene expression data analysis. Genome Biology, 19(1). https://doi.org/10.1186/s13059-017-1382-0

Zanoni, I., Ostuni, R., Marek, L. R., Barresi, S., Barbalat, R., Barton, G. M., Granucci, F., & Kagan, J. C. (2011). CD14 controls the LPS-induced endocytosis of toll-like receptor 4. Cell, 147(4), 868–880. https://doi.org/10.1016/j.cell.2011.09.051

