## Supplementary figures for "Whole-transcriptomic profiling of human cerebral cortex tissues reveals microglia-associated molecular subtypes"

### Supplemental material

#### Figure S1

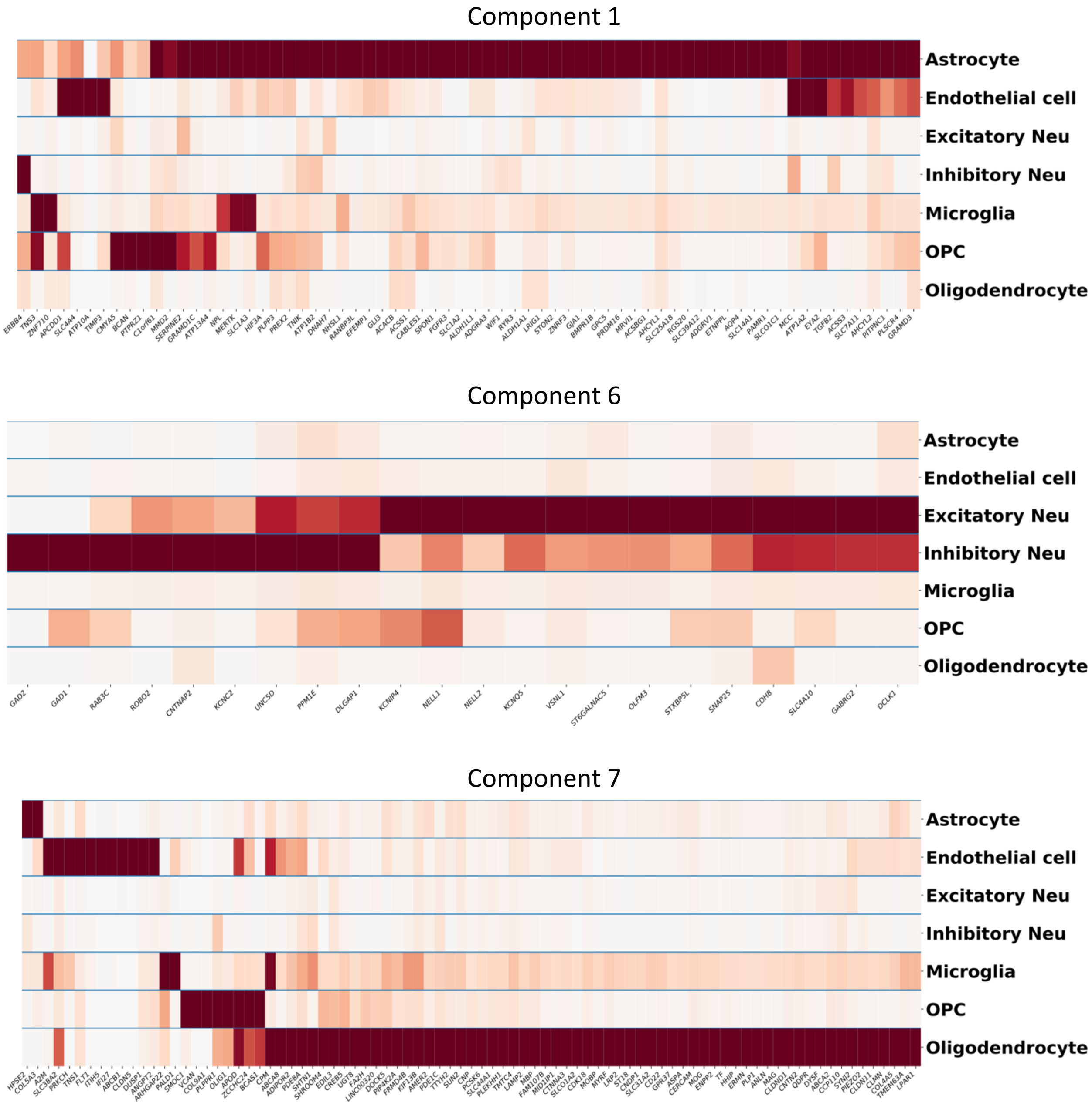


Cell type specificity of genes enriched in NMF component 1, 6, and 7.

### Figure S2

| GO.ID | Term | Hit | FDR | Module |
|---|---|---|---|---|
| GO:0050804 | modulation of chemical synaptic transmis... | 15 | 0.01376165 | purple |
| GO:0099177 | regulation of trans-synaptic signaling | 15 | 0.01376165 | |
| GO:0007610 | behavior | 15 | 0.035306857 | |
| GO:0007268 | chemical synaptic transmission | 17 | 0.035306857 | |
| GO:0098916 | anterograde trans-synaptic signaling | 17 | 0.035306857 | |
| GO:0099537 | trans-synaptic signaling | 17 | 0.035306857 | |
| GO:0099536 | synaptic signaling | 17 | 0.035306857 | |
| GO:0048167 | regulation of synaptic plasticity | 9 | 0.04072325 | |
| GO:0050806 | positive regulation of synaptic transmis... | 8 | 0.045446636 | |
| GO:0060078 | regulation of postsynaptic membrane pote... | 7 | 0.045446636 | |
| GO:0050877 | nervous system process | 28 | 0.00365105 | greenyellow |
| GO:0034220 | ion transmembrane transport | 32 | 0.005617 | |
| GO:0086010 | membrane depolarization during action po... | 6 | 0.00921188 | |
| GO:0006812 | cation transport | 30 | 0.00921188 | |
| GO:0030001 | metal ion transport | 25 | 0.00921188 | |
| GO:0098655 | cation transmembrane transport | 25 | 0.00982975 | |
| GO:0042391 | regulation of membrane potential | 17 | 0.00982975 | |
| GO:0043269 | regulation of ion transport | 21 | 0.00982975 | |
| GO:0098660 | inorganic ion transmembrane transport | 24 | 0.0101106 | |
| GO:0034765 | regulation of ion transmembrane transpor... | 17 | 0.015647357 | |
| GO:0050776 | regulation of immune response | 39 | 2.30E-22 | cyan |
| GO:0002274 | myeloid leukocyte activation | 35 | 6.74E-22 | |
| GO:0002443 | leukocyte mediated immunity | 36 | 1.46E-21 | |
| GO:0002684 | positive regulation of immune system pro... | 39 | 1.46E-21 | |
| GO:0002682 | regulation of immune system process | 45 | 1.68E-21 | |
| GO:0045321 | leukocyte activation | 41 | 9.13E-21 | |
| GO:0001775 | cell activation | 43 | 9.13E-21 | |
| GO:0002252 | immune effector process | 40 | 6.05E-20 | |
| GO:0002250 | adaptive immune response | 24 | 1.68E-19 | |
| GO:0006952 | defense response | 42 | 1.97E-19 | |
| GO:0070647 | protein modification by small protein co... | 21 | 0.004104731 | midnightblue |
| GO:0045892 | negative regulation of transcription, DN... | 19 | 0.014526724 | |
| GO:1903507 | negative regulation of nucleic acid-temp... | 19 | 0.023128824 | |
| GO:1902679 | negative regulation of RNA biosynthetic ... | 19 | 0.023128824 | |
| GO:0051253 | negative regulation of RNA metabolic pro... | 19 | 0.044413488 | |
| GO:0043543 | protein acylation | 8 | 0.044680682 | |
| GO:0006473 | protein acetylation | 7 | 0.057340208 | |
| GO:0032446 | protein modification by small protein co... | 16 | 0.065340612 | |
| GO:0016567 | protein ubiquitination | 15 | 0.067404 | |
| GO:2000113 | negative regulation of cellular macromol... | 19 | 0.073453077 | |
| GO:0072359 | circulatory system development | 7 | 1 | |
| GO:0048870 | cell motility | 8 | 1 | |
| GO:0051674 | localization of cell | 8 | 1 | |

| | | | | |
|---|---|---|---|---|
| GO:0048646 | anatomical structure formation involved ... | 6 | 1 | lightcyan |
| GO:0051726 | regulation of cell cycle | 7 | 1 | |
| GO:0016477 | cell migration | 7 | 1 | |
| GO:0044092 | negative regulation of molecular functio... | 6 | 1 | |
| GO:0022402 | cell cycle process | 7 | 1 | |
| GO:0006952 | defense response | 6 | 1 | |
| GO:0045893 | positive regulation of transcription, DN... | 6 | 1 | |
| GO:0022610 | biological adhesion | 24 | 4.04E-07 | lightgreen |
| GO:0007155 | cell adhesion | 23 | 1.26E-06 | |
| GO:0003158 | endothelium development | 8 | 7.72E-05 | |
| GO:0016477 | cell migration | 21 | 7.72E-05 | |
| GO:0048870 | cell motility | 21 | 0.000189574 | |
| GO:0051674 | localization of cell | 21 | 0.000189574 | |
| GO:0045446 | endothelial cell differentiation | 7 | 0.000205957 | |
| GO:0001525 | angiogenesis | 12 | 0.000234893 | |
| GO:0048646 | anatomical structure formation involved ... | 17 | 0.000234893 | |
| GO:0030334 | regulation of cell migration | 15 | 0.000631913 | |
| GO:0043436 | oxoacid metabolic process | 7 | 1 | lightyellow |
| GO:0006082 | organic acid metabolic process | 7 | 1 | |
| GO:0006396 | RNA processing | 7 | 1 | |
| GO:0019752 | carboxylic acid metabolic process | 6 | 1 | |
| GO:0006978 | DNA damage response, signal transduction... | 6 | 0.010030357 | turquoise |
| GO:0042772 | DNA damage response, signal transduction... | 6 | 0.0134808 | |
| GO:0006260 | DNA replication | 24 | 0.023805381 | |
| GO:0036297 | interstrand cross-link repair | 8 | 0.110333929 | |
| GO:0006259 | DNA metabolic process | 55 | 0.265871333 | |
| GO:0006261 | DNA-dependent DNA replication | 13 | 0.412498438 | |
| GO:0006302 | double-strand break repair | 17 | 0.86173 | |
| GO:0042770 | signal transduction in response to DNA d... | 12 | 0.86173 | |
| GO:0002250 | adaptive immune response | 17 | 1 | |
| GO:0030330 | DNA damage response, signal transduction... | 10 | 1 | |
| GO:0051591 | response to cAMP | 6 | 0.000584168 | darkred |
| GO:1902107 | positive regulation of leukocyte differe... | 6 | 0.00151659 | |
| GO:0046683 | response to organophosphorus | 6 | 0.00151659 | |
| GO:0042127 | regulation of cell proliferation | 17 | 0.002073969 | |
| GO:0010648 | negative regulation of cell communicatio... | 17 | 0.002073969 | |
| GO:0023057 | negative regulation of signaling | 17 | 0.002073969 | |
| GO:0014074 | response to purine-containing compound | 6 | 0.002209353 | |
| GO:0007623 | circadian rhythm | 7 | 0.002780415 | |
| GO:1903708 | positive regulation of hemopoiesis | 6 | 0.003456615 | |
| GO:0051592 | response to calcium ion | 6 | 0.004563813 | |
| GO:0006119 | oxidative phosphorylation | 14 | 1.97E-13 | |
| GO:0046034 | ATP metabolic process | 16 | 1.85E-12 | |
| GO:0009205 | purine ribonucleoside triphosphate metab... | 16 | 3.61E-12 | |

| | | | | |
|---|---|---|---|---|
| GO:0009126 | purine nucleoside monophosphate metaboli... | 16 | 3.61E-12 | darkturquoise |
| GO:0009167 | purine ribonucleoside monophosphate meta... | 16 | 3.61E-12 | |
| GO:0009144 | purine nucleoside triphosphate metabolic... | 16 | 3.61E-12 | |
| GO:0009199 | ribonucleoside triphosphate metabolic pr... | 16 | 3.61E-12 | |
| GO:0009161 | ribonucleoside monophosphate metabolic p... | 16 | 5.83E-12 | |
| GO:0009141 | nucleoside triphosphate metabolic proces... | 16 | 6.05E-12 | |
| GO:0009123 | nucleoside monophosphate metabolic proce... | 16 | 8.99E-12 | |
| GO:0060337 | type I interferon signaling pathway | 21 | <1E-25 | darkgrey |
| GO:0071357 | cellular response to type I interferon | 21 | <1E-25 | |
| GO:0034340 | response to type I interferon | 21 | <1E-25 | |
| GO:0045087 | innate immune response | 29 | <1E-25 | |
| GO:0006952 | defense response | 32 | <1E-25 | |
| GO:0019221 | cytokine-mediated signaling pathway | 26 | 7.86E-25 | |
| GO:0051607 | defense response to virus | 18 | 4.86E-22 | |
| GO:0071345 | cellular response to cytokine stimulus | 26 | 1.08E-20 | |
| GO:0098542 | defense response to other organism | 19 | 1.35E-20 | |
| GO:0034097 | response to cytokine | 26 | 5.52E-20 | |
| GO:0044282 | small molecule catabolic process | 49 | 2.25E-13 | blue |
| GO:0016054 | organic acid catabolic process | 39 | 2.25E-13 | |
| GO:0046395 | carboxylic acid catabolic process | 39 | 2.25E-13 | |
| GO:0019752 | carboxylic acid metabolic process | 74 | 1.80E-10 | |
| GO:0006082 | organic acid metabolic process | 78 | 1.80E-10 | |
| GO:0043436 | oxoacid metabolic process | 76 | 7.86E-10 | |
| GO:0055114 | oxidation-reduction process | 68 | 1.60E-09 | |
| GO:0003008 | system process | 85 | 4.21E-09 | |
| GO:0007423 | sensory organ development | 42 | 9.99E-09 | |
| GO:1901605 | alpha-amino acid metabolic process | 28 | 2.25E-08 | |
| GO:0001568 | blood vessel development | 62 | 1.01E-17 | brown |
| GO:0016477 | cell migration | 93 | 1.01E-17 | |
| GO:0072358 | cardiovascular system development | 64 | 1.01E-17 | |
| GO:0048514 | blood vessel morphogenesis | 57 | 1.01E-17 | |
| GO:0001944 | vasculature development | 63 | 1.12E-17 | |
| GO:0048870 | cell motility | 97 | 2.01E-17 | |
| GO:0051674 | localization of cell | 97 | 2.01E-17 | |
| GO:0001525 | angiogenesis | 51 | 8.43E-17 | |
| GO:0035239 | tube morphogenesis | 65 | 4.44E-16 | |
| GO:0048646 | anatomical structure formation involved ... | 74 | 1.47E-15 | |
| GO:0043436 | oxoacid metabolic process | 36 | 0.007676567 | yellow |
| GO:0006082 | organic acid metabolic process | 36 | 0.007676567 | |
| GO:0019752 | carboxylic acid metabolic process | 34 | 0.007676567 | |
| GO:0017144 | drug metabolic process | 27 | 0.0269616 | |
| GO:0032787 | monocarboxylic acid metabolic process | 21 | 0.0758295 | |
| GO:0055114 | oxidation-reduction process | 29 | 0.079440429 | |
| GO:0006767 | water-soluble vitamin metabolic process | 7 | 0.106098889 | |
| GO:0006520 | cellular amino acid metabolic process | 16 | 0.106098889 | |

| | | | | |
|---|---|---|---|---|
| GO:0006629 | lipid metabolic process | 34 | 0.16851 | |
| GO:0005975 | carbohydrate metabolic process | 20 | 0.16851 | |
| GO:0060271 | cilium assembly | 12 | 0.2331055 | green |
| GO:0044782 | cilium organization | 12 | 0.2331055 | |
| GO:0003002 | regionalization | 8 | 1 | |
| GO:0120031 | plasma membrane bounded cell projection ... | 13 | 1 | |
| GO:0030031 | cell projection assembly | 13 | 1 | |
| GO:0006325 | chromatin organization | 14 | 1 | |
| GO:0007389 | pattern specification process | 8 | 1 | |
| GO:0031503 | protein-containing complex localization | 8 | 1 | |
| GO:0070925 | organelle assembly | 14 | 1 | |
| GO:0008380 | RNA splicing | 10 | 1 | |
| GO:0042552 | myelination | 21 | 6.74E-11 | red |
| GO:0007272 | ensheathment of neurons | 21 | 6.74E-11 | |
| GO:0008366 | axon ensheathment | 21 | 6.74E-11 | |
| GO:0048709 | oligodendrocyte differentiation | 18 | 2.25E-10 | |
| GO:0042063 | gliogenesis | 25 | 1.06E-08 | |
| GO:0010001 | glial cell differentiation | 22 | 1.59E-08 | |
| GO:0007417 | central nervous system development | 45 | 7.86E-07 | |
| GO:0014003 | oligodendrocyte development | 11 | 8.43E-07 | |
| GO:0022010 | central nervous system myelination | 8 | 1.53E-06 | |
| GO:0032291 | axon ensheathment in central nervous sys... | 8 | 1.53E-06 | |
| GO:0006457 | protein folding | 27 | 1.35E-16 | black |
| GO:0006986 | response to unfolded protein | 21 | 2.25E-12 | |
| GO:0035966 | response to topologically incorrect prot... | 22 | 2.25E-12 | |
| GO:0061077 | chaperone-mediated protein folding | 14 | 7.30E-12 | |
| GO:0009408 | response to heat | 19 | 4.16E-11 | |
| GO:0009266 | response to temperature stimulus | 20 | 6.65E-10 | |
| GO:1900034 | regulation of cellular response to heat | 14 | 1.36E-09 | |
| GO:0034605 | cellular response to heat | 16 | 1.61E-09 | |
| GO:0006458 | 'de novo' protein folding | 9 | 5.37E-07 | |
| GO:0090084 | negative regulation of inclusion body as... | 6 | 7.86E-07 | |
| GO:0002252 | immune effector process | 46 | 1.17E-09 | pink |
| GO:0001775 | cell activation | 46 | 5.48E-08 | |
| GO:0006952 | defense response | 46 | 1.31E-07 | |
| GO:0034097 | response to cytokine | 41 | 6.09E-07 | |
| GO:0071345 | cellular response to cytokine stimulus | 39 | 6.09E-07 | |
| GO:0002443 | leukocyte mediated immunity | 32 | 6.91E-07 | |
| GO:0072358 | cardiovascular system development | 31 | 9.36E-07 | |
| GO:0045321 | leukocyte activation | 40 | 9.48E-07 | |
| GO:0001944 | vasculature development | 30 | 1.79E-06 | |
| GO:0006954 | inflammatory response | 25 | 5.17E-06 | |

Top 10 most enriched Gene Ontology (biological processes) terms for each of the modules identified with WGCNA.

### Figure S3

a

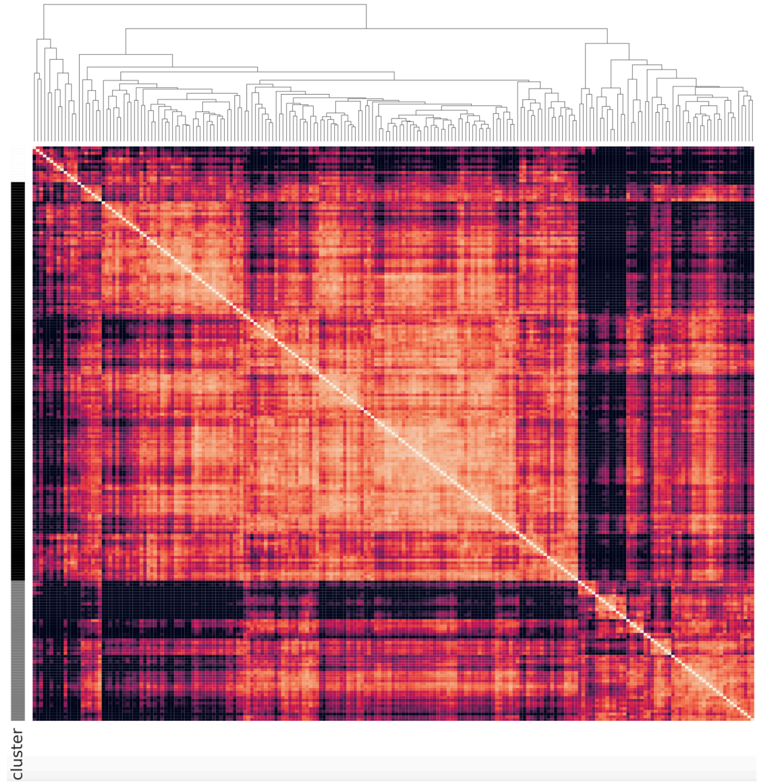


b

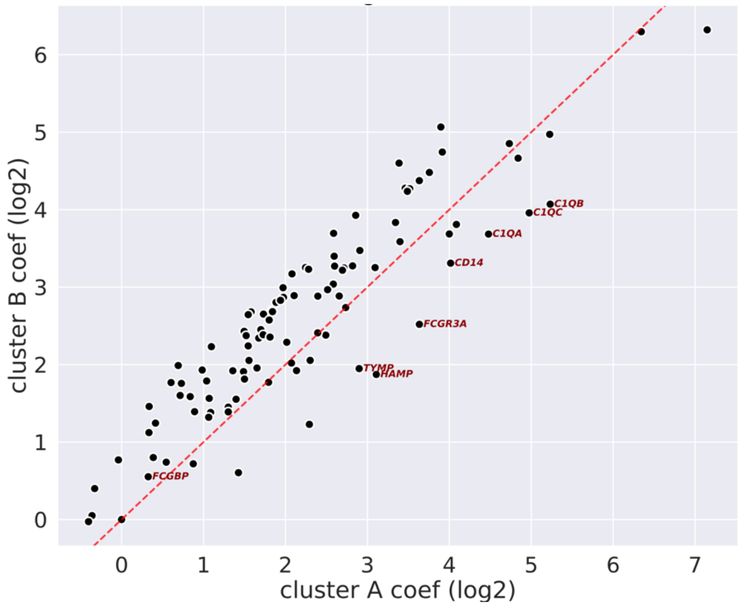


Hierarchical clustering of GTEx Frontal Cortex samples based on the expression levels of microglia genes (a). Grey and black bars represent ClusterA and ClusterB, respectively. C1 complement system components as well FCGR3A and CD14 are more highly expressed in ClusterA (b).

Figure S4

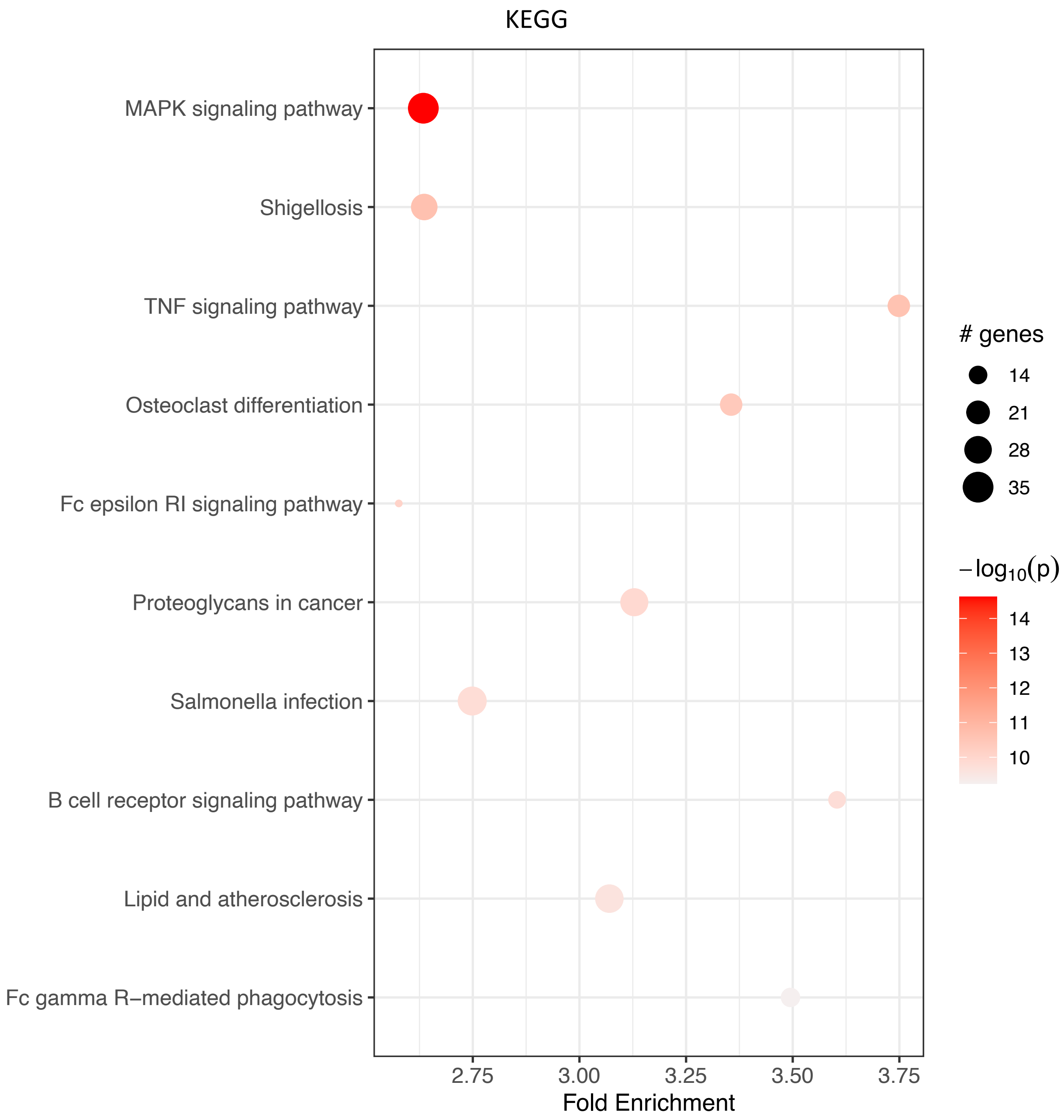
KEGG
MAPK signaling pathway
Shigellosis
TNF signaling pathway
Osteoclast differentiation
Fc epsilon RI signaling pathway
Proteoglycans in cancer
Salmonella infection
B cell receptor signaling pathway
Lipid and atherosclerosis
Fc gamma R–mediated phagocytosis
2.75
3.00
3.25
3.50
3.75
Fold Enrichment
### genes
14
21
28
35
$-\log_{10}(p)$
14
13
12
11
10

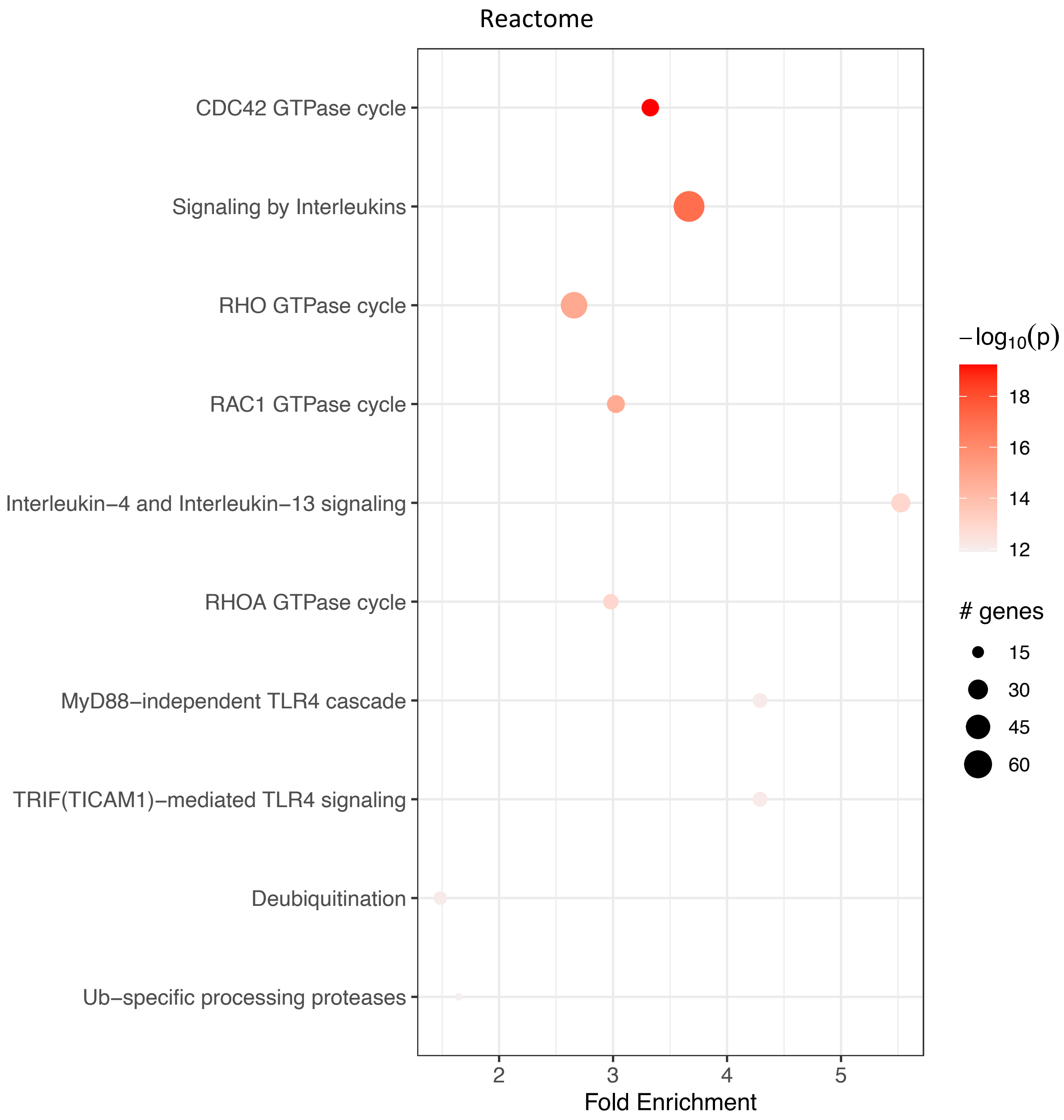


KEGG and Reactome pathway enrichment for genes differentially expressed between two subtypes of microglia.

### Table S1

| ROSMAP | | MSBB | |
|---|---|---|---|
| Gene Name | ClusterB Coef. - ClusterA Coef. | Gene Name | ClusterB Coef. - ClusterA Coef. |
| FCGR3A | -1.032411114 | FCGBP | -1.281003683 |
| HAMP | -1.021032471 | FCGR3A | -0.851004638 |
| C1QB | -0.967937193 | C1QB | -0.831837046 |
| FCGBP | -0.89494333 | C1QC | -0.668336261 |
| C1QC | -0.881667972 | C1QA | -0.618116524 |
| CD14 | -0.822152458 | CD14 | -0.574400264 |
| C1QA | -0.794094169 | STAB1 | -0.337999111 |
| TYMP | -0.753580116 | TYMP | -0.32948512 |
| STAB1 | -0.528159536 | ITGB2 | -0.270432883 |
| ITGB2 | -0.513425107 | C3 | -0.222386138 |

Comparison of top10 differential genes between ClusterA and ClusterB in the ROSMAP and MSBB cohorts. Genes shared in both cohorts are represented in red.
